# WhiB6 is required for the secretion-dependent regulation of ESX-1 substrates in pathogenic mycobacteria

**DOI:** 10.1101/297440

**Authors:** Abdallah M. Abdallah, E.M. Weerdenburg, Qingtian Guan, R. Ummels, S. Borggreve, S.A. Adroub, Tareq B. Malas, Raeece Naeem, Huoming Zhang, T.D. Otto, W. Bitter, A. Pain

**Affiliations:** Pathogen Genomics Laboratory, BESE Division, King Abdullah University of Science and Technology (KAUST), Thuwal-Jeddah, Kingdom of Saudi Arabia; Department of Medical Microbiology and Infection Control, VU University Medical Center, Amsterdam, The Netherlands; Bioscience Core Laboratory, King Abdullah University of Science and Technology (KAUST), Thuwal-Jeddah, Kingdom of Saudi Arabia; Pathogen Genomics, The Wellcome Trust Sanger Institute, Hinxton, Cambridge, United Kingdom

**Author notes:** Joint authors (Abdallah M. Abdallah, E.M. Weerdenburg). Joint authors (W. Bitter, A. Pain). Corresponding authors Abdallah M. Abdallah, Wilbert Bitter, Arnab Pain.

## Abstract

The mycobacterial type VII secretion system ESX-1 is responsible for the secretion of a number of proteins that play important roles during host infection. The regulation of the expression of secreted proteins is often essential to establish successful infection. Using transcriptome sequencing, we found that the abrogation of ESX-1 function in *Mycobacterium marinum* leads to a pronounced increase in gene expression levels of the *espA* operon during the infection of macrophages, suggesting an important role in ESX-1-mediated virulence during the early phase of infection. In addition, the disruption of ESX-1-mediated protein secretion also leads to a specific down-regulation of the ESX-1 substrates, but not of the structural components of this system, during growth in culture medium. This effect is observed in both *M. marinum* and *M. tuberculosis*. We established that down-regulation of ESX-1 substrates is the result of a regulatory process that is influenced by the putative transcriptional regulator *whib6*, which is located adjacent to the *esx-1* locus. In addition, the overexpression of the ESX-1-associated PE35/PPE68 protein pair resulted in a significantly increased secretion of the ESX-1 substrate EsxA, demonstrating a functional link between these proteins. Taken together, these data show that WhiB6 is required for the secretion-dependent regulation of ESX-1 substrates and that ESX-1 substrates are regulated independently from the structural components, both during infection and as a result of active secretion.

## Introduction

Mycobacteria use several different type VII secretion systems (T7S) to transport proteins across their thick and waxy cell envelopes. One of these T7S systems, ESX-1, is responsible for the transport of a number of important virulence factors. Disruption of the *esx-1* gene cluster severely reduces the virulence of *Mycobacterium tuberculosis* [1], whereas restoration of *esx-1* in the *Mycobacterium* bovis-derived vaccine strain BCG, which lacks part of the *esx-1* region due to continuous passaging, leads to increased virulence [2]. Many studies have attempted to elucidate the function of ESX-1 substrates in virulence. In the case of pathogenic mycobacteria, such as *M. tuberculosis* and the fish pathogen *Mycobacterium marinum*, ESX-1 is responsible for the translocation of the bacteria from the phagolysosomal compartments to the cytosols of macrophages [3,4]. This translocation activity has been attributed to the ability of the secreted protein EsxA (also called ESAT-6) to lyse membranes [5,6]. Interestingly, a closely related homologue of this protein is also secreted by non-pathogenic and non-translocating mycobacteria such as *Mycobacterium smegmatis*. A recent report indicated that, although the EsxA proteins of *M. smegmatis* and *M. tuberculosis* are highly homologous, the membrane lysis potentials of these proteins are different [7]. In *M. smegmatis*, ESX-1 is involved in a completely different process, *i.e*., conjugative DNA transfer [8]. The proposed functions of ESX-1 in pathogenic mycobacterial species include host cell entry and intercellular spread [9].

The ESX-1 substrates identified to date are mostly encoded by genes of the *esx-1* locus, such as EsxA, EsxB, EspE and EspB. The exceptions are EspA and EspC [10,11], which are both part of the *espA* operon, which is located elsewhere in the genome; however, these genes are homologous to the *espE* and *espF* genes, respectively, which belong to the *esx-1* locus. A peculiar characteristic of ESX-1 substrates is that these substrates are mutually dependent, *i.e*., the secretion of each of these substrates is dependent on the secretion of the other substrates [10]. The secreted ESX proteins contain a conserved WxG amino acid motif located between two α-helices [12]. Recently, an additional conserved secretion signal, present in all secreted protein pairs, was identified. This C-terminal YxxxD/E motif can target proteins for secretion, but does not determine the specificity for a particular type VII system [13]. Therefore, it remains difficult to bioinformatically predict novel ESX-1 substrates.

To establish successful infection, mycobacteria need regulatory mechanisms to express the right proteins at the right time. In different environments, mycobacteria require specific transcriptional responses to successfully respond to the stress conditions encountered. During the first stages of infection, ESX-1-mediated protein secretion is one of the most important virulence mechanisms of pathogenic mycobacteria [4,9,10,14,15]. Consequently, the tight transcriptional regulation of *esx-1* and the associated genes is required. The transcriptional regulator PhoP of the two-component system PhoPR positively regulates the transcription of many esx-1-associated genes, including genes in the *espA* operon [16,17]. It has been proposed that PhoP regulation is dependent on environmental pH [18], which could indicate that the acidic environment of the phagosome induces *esx-1* gene transcription via PhoP, leading to bacterial escape from this compartment. Other studies have shown that the *espA* operon is, in addition to PhoP, also regulated by the transcription factors EspR and MprAB and the repressors CRP and Lsr2, indicating that tight regulation of this operon is essential and, furthermore, suggesting that the *espA* operon may be regulated in a manner distinct from the regulation of other ESX-1 substrates [19–22].

Since ESX-1 is crucial for virulence, inactivation of this secretion system would be expected to have a large impact on gene regulation processes in mycobacteria. Here, we apply RNA-seq and quantitative proteomics to determine the gene expression and proteomic profiles of the pathogenic mycobacteria *M. marinum* and *M. tuberculosis* in the absence of a functional ESX-1 secretion system. During short-term infection of macrophages, we observed highly increased transcript levels of the *espA* operon. In contrast, during in vitro growth in culture medium, transcription of most ESX-1 substrates and some putative new substrates was seen to be decreased. Based on these gene transcription levels, we confirmed a regulatory role for the putative transcriptional regulator WhiB6 in the gene expression of ESX-1 substrates.

## Results

### Global features of the M. marinum esx-1 mutant transcriptome and proteome

To investigate the effect of ESX-1 disruption on gene expression and protein production, RNA and protein were extracted from three independent exponential phase cultures of the *M. marinum* E11 strain and the isogenic esx-1-mutant during growth in 7H9 culture medium to characterize the transcriptome and proteome. Using transcriptomics (RNA-seq) and mass spectrometry (MS)-based proteomics with isobaric labelling for quantification, we captured the expression dynamics of the transcripts and proteomes of the *esx-1* mutant. Data quality was assessed using Euclidean distance matrices for RNA (Figure S1) and principal component analysis (PCA) for protein (Figure S2), which demonstrated high levels of reproducibility between biological replicates. After filtering (see Materials and Methods for details), a total of 823 genes were identified as being differentially expressed (DE) as messenger RNA, of which 525 were classified as down-regulated and 298 as up-regulated (Figure 1A, Table S1). To determine parallel changes in protein levels, 1,657 proteins were identified by the presence of 2 or more peptides, of which 576 proteins passed our filter and we classified them as DE. Of these, 412 proteins were found to be down-regulated and 164 were up-regulated (Table S2), and 482 protein-coding genes were shared and were identified in both the RNA-seq and quantitative proteomic datasets (Figure 1C).

**Fig 1.**
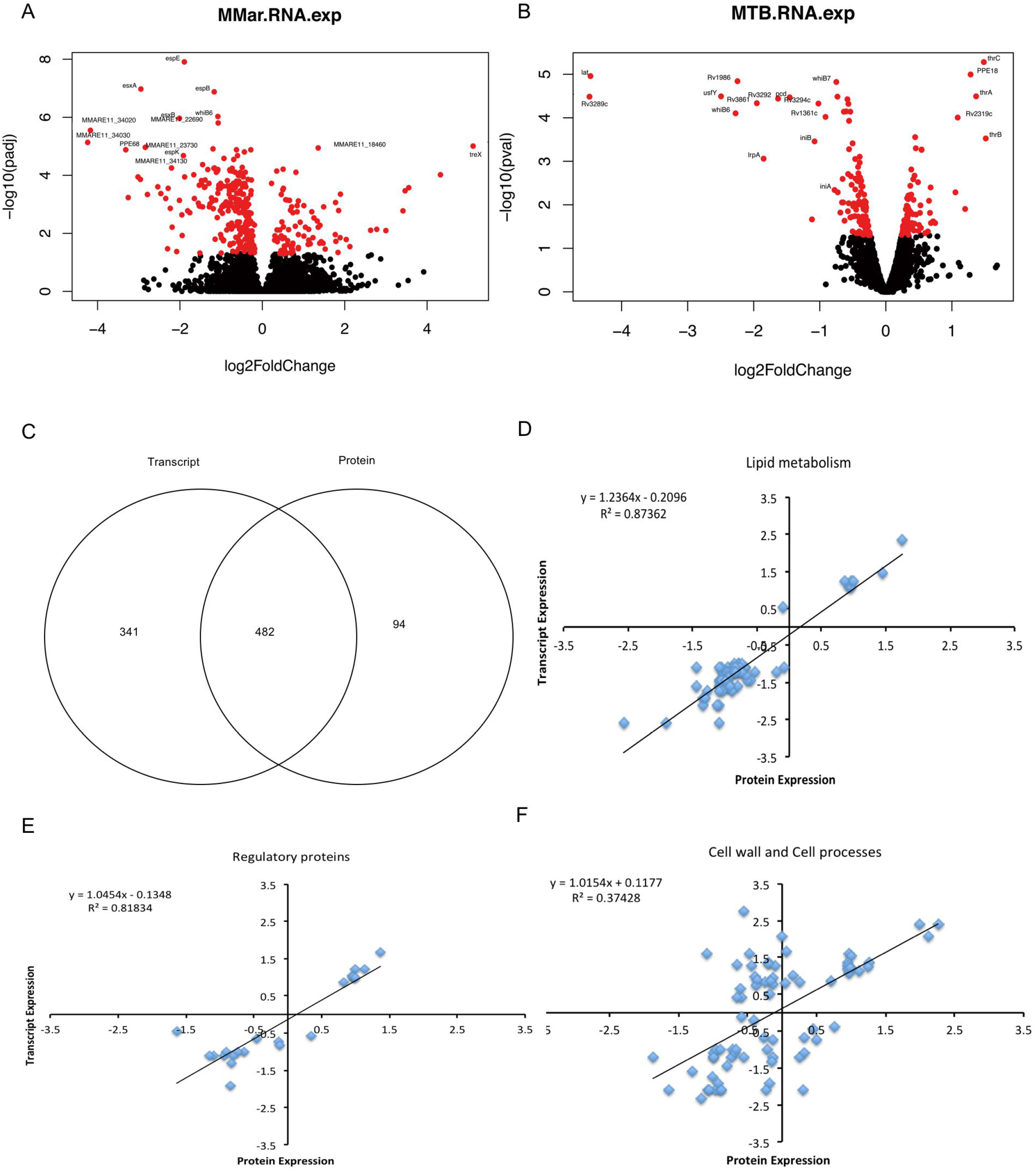
Global Features of the Transcriptomes and Proteomes of the *M. marinum* and *M. tuberculosis* Esx-1 Mutant Strains. Volcano plots obtained from RNA-seq analysis of the wild-type *M. marinum* E11 strain vs. the *eccCb_1_* transposon mutant (A) and of *M. tuberculosis* mc^2^6020 vs. the ESX-1 mutant strain (B). Each dot indicates the expression value of a gene. Red dots indicate statistical significance (q < 0.05), and black dots indicate a lack of statistical significance. Selected genes that are most down- or up-regulated in the *ESX-1* mutant strains are highlighted. (C) Venn diagram of the number of differentially expressed transcripts and proteins quantified using RNA-seq and quantitative proteomics, respectively. Scatterplots of the relationship between differentially expressed genes of *M. marinum eccCb_1_* transposon mutant and those of the isogenic wild-type strain E11, quantified in both data sets and classified into the following categories: (D) lipid metabolism, (E) regulatory proteins and (F) cell wall and cell process. Scatterplots and bar chart show the rectilinear equation and the Pearson correlation coefficient (R^2^).

The degree of global correlation between the gene expression and protein abundance scores among the shared genes was relatively low (Figure S3A), which has also been noted in other bacterial studies [23]. However, within certain classes of *M. marinum* functional categories (http://mycobrowser.epfl.ch/marinolist.html), the degree of correlation was much higher than that in other classes, with R^2^ exceeding 0.8 for the categories such as lipid metabolism (Figure 1D), regulation (Figure 1E) and cell wall and cell processes (Figure 1F). Of the DE genes at the RNA and protein levels, 28% were in the intermediary metabolism and respiration category, 18% were in the cell wall and cell process category, 15% were in the information pathways category and 14% were in the lipid metabolism category (Figure S4).

Transcriptional profiling analysis of the double auxotrophic *M. tuberculosis* mc^2^6020 mutant strains [24] and their isogenic *esx-1* mutants during growth was carried out to identify genes for which expression was dependent on ESX-1 disruption (Figure 1B, Table S3). For this species, the same trends could be identified as for *M. marinum*.

### Major effects of ESX1 mutation on genes encoding ESX-1 substrates and biosynthetic pathways

Analysis of differential expression in *M. marinum* identified changes in genes involved in a variety of cellular processes (Figure 2), although a majority of the most differentially regulated genes were associated with cell wall and cell processes and lipid metabolism. We noted that a substantial number of esx-1-associated genes were down-regulated in the mutant strains during growth in culture medium, including 11 genes that were located within or directly adjacent to the *esx-1* gene cluster. Among these down-regulated genes were those coding for known ESX-1 substrates, such as EsxA, EsxB, EspE and EspB. Remarkably, mRNA levels of core components of the ESX-1 secretion system, *i.e*., those encoding members of the type VII secretion complex, such as EccB_1_, EccD_1_, EccE_1_ and MycP_1_, remained unchanged, even though the corresponding genes are interspersed with genes encoding ESX-1 substrates. In contrast to the mRNA levels, we noted a strong increase in the protein levels of EsxA and EsxB, probably reflecting the accumulation of these proteins in the cell due to the secretion defect (Figure 2). Our data also indicate a significant effect of *esx-1* disruption on genes associated with lipid metabolism (Figure 2), including genes associated with the synthesis of mycolic acids. Strong down-regulation was observed at the mRNA and protein levels for several polyketide synthases, including genes involved in phthiocerol dimycocerosate synthesis and mycolic acid biosynthesis, such as *umaA, mmaA3, accD5, accD6*, and *pks15/1*, which encode components of the lipid biosynthesis pathway (Figure 2, Table S1, S2). The changes observed in *esx-1* and lipid-metabolism-associated genes at the mRNA and protein levels were not unexpected; it has been reported previously that ESX-1-dependent protein secretion and mycolic acid synthesis are critically linked [25]. However, we also noted a surprisingly broad impact of ESX-1 mutation on major biosynthetic pathways, including ribosomal protein synthesis and DNA biosynthesis (Table S1, S2). Down-regulation was observed at the mRNA and protein levels for several genes encoding ribosomal proteins and DNA gyrase and a ribonucleotide-diphosphate reductase, which are components of protein and DNA biosynthesis, respectively. We also identified changes at both the mRNA and protein levels in genes involved in general stress response (*grpE*, *dnaK*, *groES*, *groEL1*), genes involved in stress response regulation (*sigA*, *sigB*, *devS*), members of the WhiB family (*whiB2*, *whiB4*, *whiB6*) and several PE_PGRS genes (Figure 2). A similar trend was observed for *M. tuberculosis* (Figure 2). In fact, of all the genes, the *esx-1* genes encoding the substrates EsxA, EsxB and EspK were the most significantly down-regulated in the mutant strain (Figure 2, Table S3). However, in contrast to the levels in the *M. marinum* mutant, the gene expression levels of *eccB1* and *eccD1* were also somewhat decreased in the *M. tuberculosis* mutant (Figure 2). On the other hand, the *M. tuberculosis esx-1* mutation did not seem to have a significant effect on the expression of genes involved in lipid metabolism compared to the effect seen in *M. marinum* (Figure 2, Table S1, S3). Finally, a significant number of genes that are associated with information pathways, including genes encoding ribosomal proteins, were up-regulated at the mRNA level in the *esx-1* mutant (Figure 2). Taken together, the observed changes in the transcriptome and proteome of the *esx-1* mutant reflect the role of the *esx-1* cluster employed by mycobacteria for major biochemical pathways.

**Fig 2.**
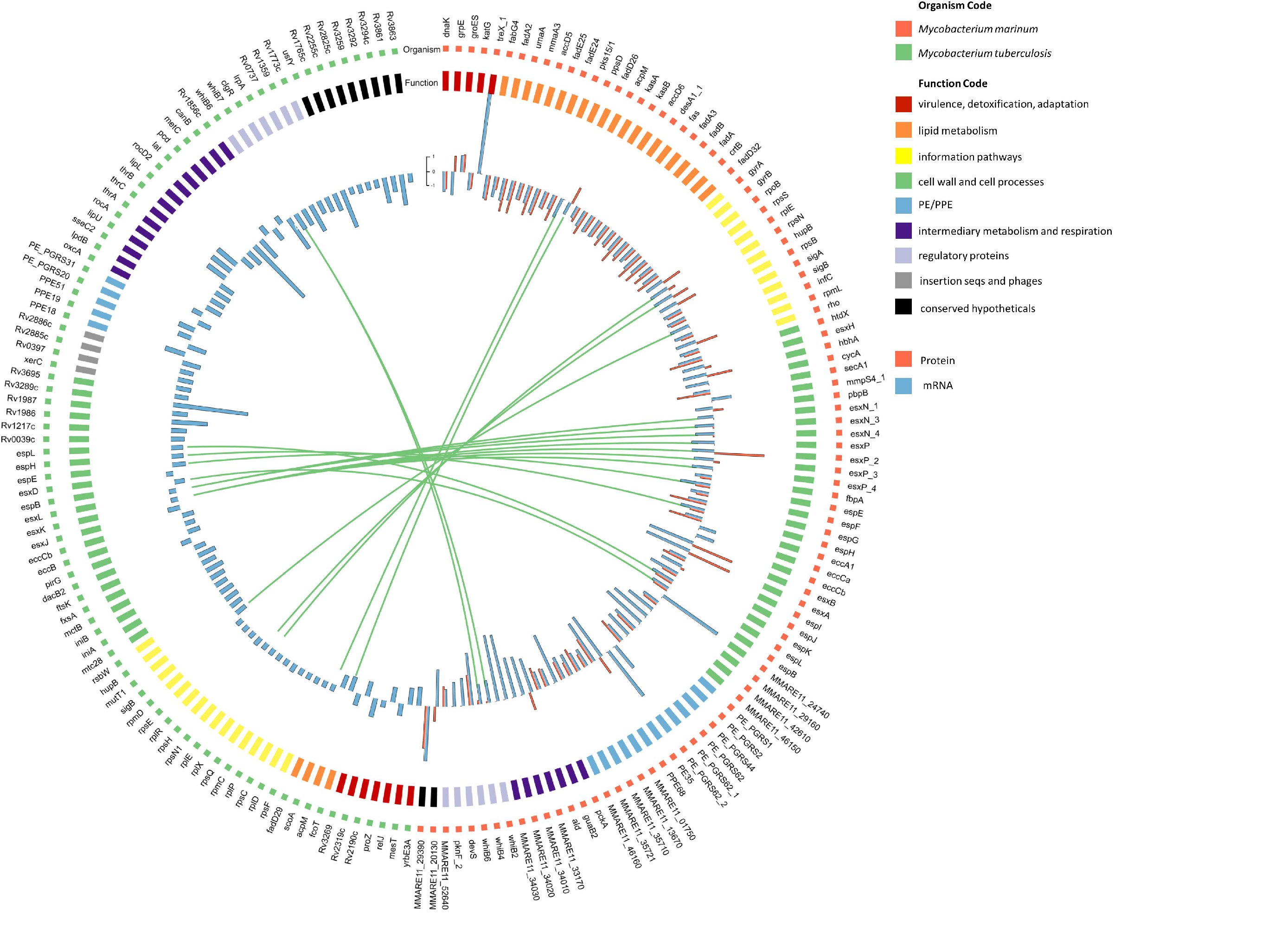
Top Differentially Expressed Genes of *M. marinum* and *M. tuberculosis*, when Grown in Culture Medium, Grouped into Broad Functional Categories. Within each group, genes are ranked in ascending order by p-value. (Red) Top 100 annotated *M. marinum* E11 genes that exhibit greatest differential expression in the *M. marinum eccCb_1_* transposon mutant compared to the isogenic wild-type strain E11 during growth in 7H9 culture medium. Bar chart of log2-fold change for individual genes (RNA, blue; protein, red; locus tags, outer). (Green) Top 100 annotated *M. tuberculosis* genes that exhibit greatest differential expression in the auxotrophic *M. tuberculosis* RD1 deletion mutant strain mc^2^6030 compared to the isogenic control strain mc^2^6020 during growth in 7H9 culture medium. Bar chart of log2-fold change for individual genes. The genes Rv3872-Rv3878 are not included as these genes are deleted in the RD1 mutant strain.

### Global transcriptional profiling of intraphagosomal M. marinum and the esx-1 mutant

We next determined the effect of ESX-1 abrogation in *M. marinum* on gene transcription during infection of primary macrophages. Using a PMA-differentiated THP-1 cell line as a model of primary macrophages, we analysed the global gene expression of the wild-type and *esx-1* mutant strains of *M. marinum* after 6 hours of infection. Wild-type mycobacteria can escape the phagosome within two hours after infection, whereas *esx-1* mutants are known to be limited to the phagosomal compartment. The intraphagosomal transcriptome of the *esx-1* mutant was compared with the intracellular transcriptome of wild-type *M. marinum*. Furthermore, these intracellular transcriptomes were also compared with the transcriptome of wild-type *M. marinum* grown in standard broth culture. We identified 720 (p<0.05) genes in the *esx-1* mutant that exhibited significant changes in expression after THP-1 infection compared to the expression levels in the wild-type strain. Of these genes, 465 were down-regulated and 255 were up-regulated (Table S4, Figure S5). Remarkably, none of the genes within the *esx-1* region were significantly differentially expressed in the esx-1-mutant compared to the wild-type strain. However, we found a specific and pronounced increase in the transcript levels of the *espA* operon in the intraphagosomal transcriptome of the *esx-1* mutant compared with the levels in the in vitro transcriptomes (Figure 3A). During growth in culture medium, the mRNA levels of *espA* did not differ between the wild-type and esx-1-deficient *M. marinum* strains, which was confirmed by quantitative RT-PCR (qRT-PCR) (Figure 3B). Therefore, these data suggest that proteins encoded by the *espA* operon, *i.e*., EspA, EspC and EspD, play an important role in ESX-1-specific processes during the first stages of macrophage infection. The *espA* operon was also induced in the wild-type bacteria inside macrophages, albeit at a lower level. Perhaps this difference exists because the wild-type bacteria are able to escape from the phagosome, whereas the *esx-1* mutants are not.

**Fig 3.**
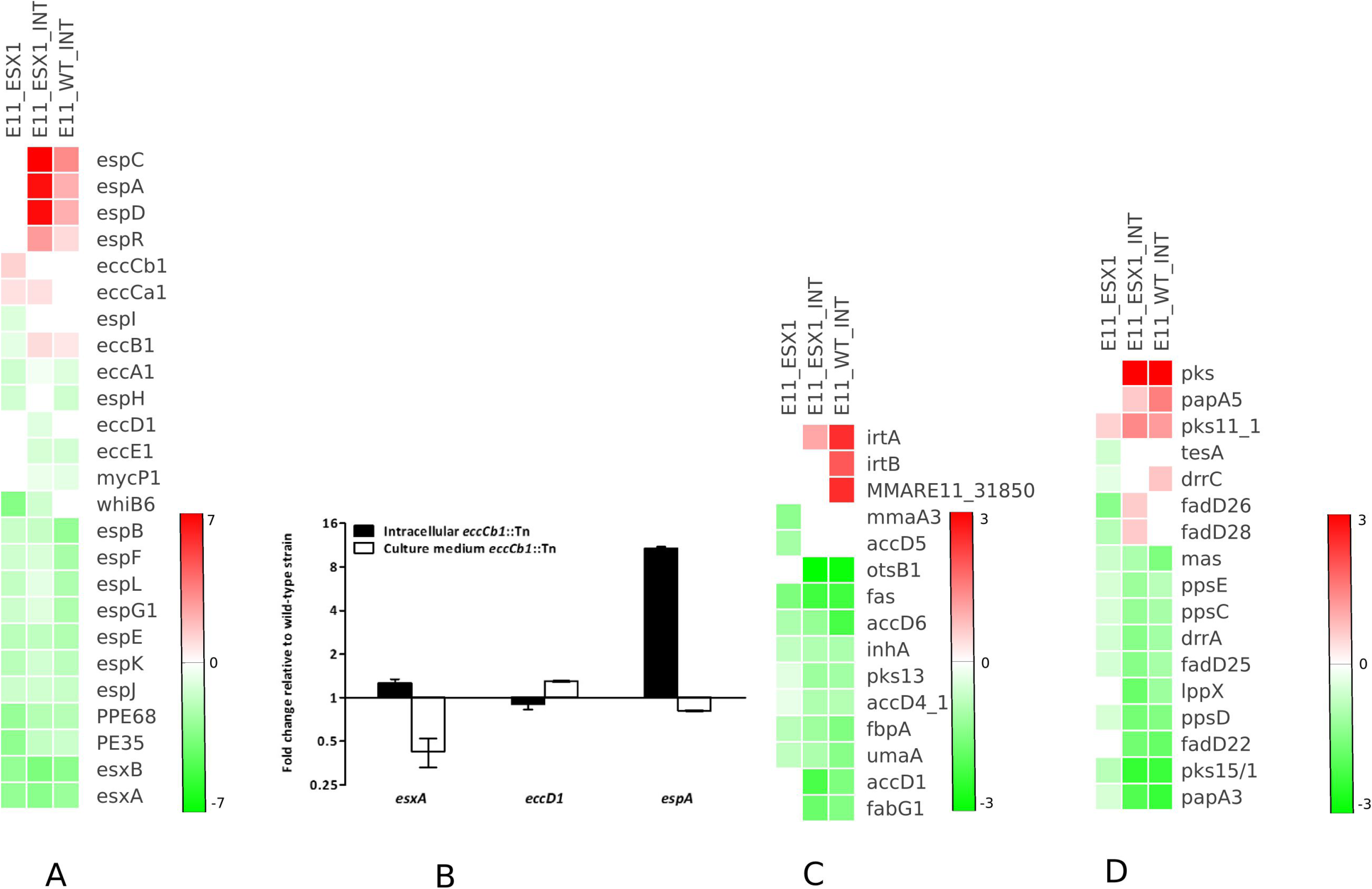
Effect of ESX-1 Disruption (*eccCb_1_* Transposon Mutant) on Gene Transcription During Infection (Indicated as ínt’) and Growth in Culture Medium in *M. marinum* Compared to that in the Wild-Type Strain E11 during Growth in 7H9 Culture Medium. (A) Relative transcript expression levels of the ESX-1 secretion system-associated genes, including the main ESX-1 locus as well as the EspR regulator and accessory factors in the EspA operon, which is encoded outside the RD-1 region. (B) Gene expression levels, as measured by qRT-PCR, were compared to those of the wild-type strain E11 grown in similar conditions. Values represent mean ± standard error of the mean of two biological replicates. (C, D) Regulation of genes associated with cell wall synthesis, including genes involved in mycolic acid synthesis (C) and PDIMs (D).

Further analysis showed that a significant number of genes that code for proteins involved in cell wall and cell processes were differentially regulated by intracellular wild-type *M. marinum* and the ESX-1-deficient strain in comparison with their counterparts grown in culture medium (Table S5, S6). *M. marinum* genes involved in mycolic acid synthesis, phthiocerol dimycocerosate (PDIM) synthesis and transport to the cell surface, such as *fabG1, accDs, ppsC, ppsD, pks11_1, pks13*, as well as genes coding for polyketide synthases and the mycolic acid methyltransferase *umaA* were differentially expressed during infection of THP-1 cells (Figure 3C, D). Furthermore, *cpsY*, a gene that encodes UDP-glucose 4-epimerases and is essential for linking peptidoglycans and mycolic acid [26], exhibited a pronounced increase in mRNA level in the intracellular *esx-1* mutant (Table S4, S5, S6, S7). We also found that many genes associated with cell division and peptidoglycan assembly, such as *ftsE*, *ftsH*, *ftsW*, *murC*, and *murG* [27,28], were down-regulated by intracellular bacteria (Table S4, S5, S6, S7).

A significant number of genes that code for proteins associated with lipid metabolism and metabolic adaptation were differentially regulated in macrophages (Figure S6A). This subset includes genes involved in fatty acid metabolism such as isocitrate lyase (*icl*), an enzyme necessary for the glyoxylate cycle and required for intracellular survival [29,30], and *pckA*, which encodes the phosphoenolpyruvate carboxykinase and is essential for mycobacterial survival in both macrophages and mice [31,32] and is involved in energy metabolism (Figure S6B), and the KstR-dependent cholesterol regulon (Figure S6C), which is involved in lipid degradation and carbon metabolism [33]. We also observed significant effects of a number of genes involved in general stress response (*groES*, *groEL1*, *hsp*, *ahp*, *dnaK*), genes involved in stress response regulation (*sigB*, *devR*, *devS*, *hspR*, *kstR*), members of the WhiB family (*whiB2*, *whiB3*, *whiB4*, *whiB*, *whiB6*, *whiB7*) and alternative sigma factors (*sigE*, *sigL*, *sigM*) in the *esx-1* mutant during infection to macrophages. This pattern is illustrated in Figure S6D and is probably associated with stressful intraphagosomal conditions.

### Different M. marinum esx-1 transposon mutants have similar gene transcription profiles

The ESX-1-deficient strain of *M. marinum* used for RNA sequencing contains a transposon in the *eccCb_1_* gene. To confirm that the observed gene transcription effects were due to a defective ESX-1 system and not due to a side effect of this particular mutation, we analysed several mutants containing transposon insertions in different genes from the *esx-1* gene cluster and compared the mRNA levels of the selected genes by qRT-PCR. Our results showed decreased transcript levels of the known ESX-1 substrate *esxA* and other *esx-1* secretion-associated (esp) genes, namely, *espL*, *espK* and *espJ*, for all tested *esx-1* mutants, whereas the transcript levels of *eccD_1_* which encodes a structural component of the ESX-1 system, did not differ from the transcript levels in wild-type *M. marinum* (Figure 4). These gene expression patterns in the *eccB_1_, eccCa_1_, eccD_1_* and *eccE_1_* transposon mutants were similar to the RNA sequencing results obtained for the *eccCb_1_* mutant. The only exception was that for the mutant containing a transposon insertion in *eccD_1_*, we observed an increase of *eccD1* transcription itself and, to a lesser extent, of the adjacent gene *espJ* (Figure 4). However, this increase was most likely due to the presence of a strong promoter on the transposon, driving the transcription of the kanamycin resistance cassette, as the measured mRNA is transcribed from sequences directly downstream of this promoter. Altogether, our results demonstrate that inactivation of the ESX-1 secretion system leads to down-regulation of the transcription of ESX-1 substrates and associated proteins.

**Fig 4.**
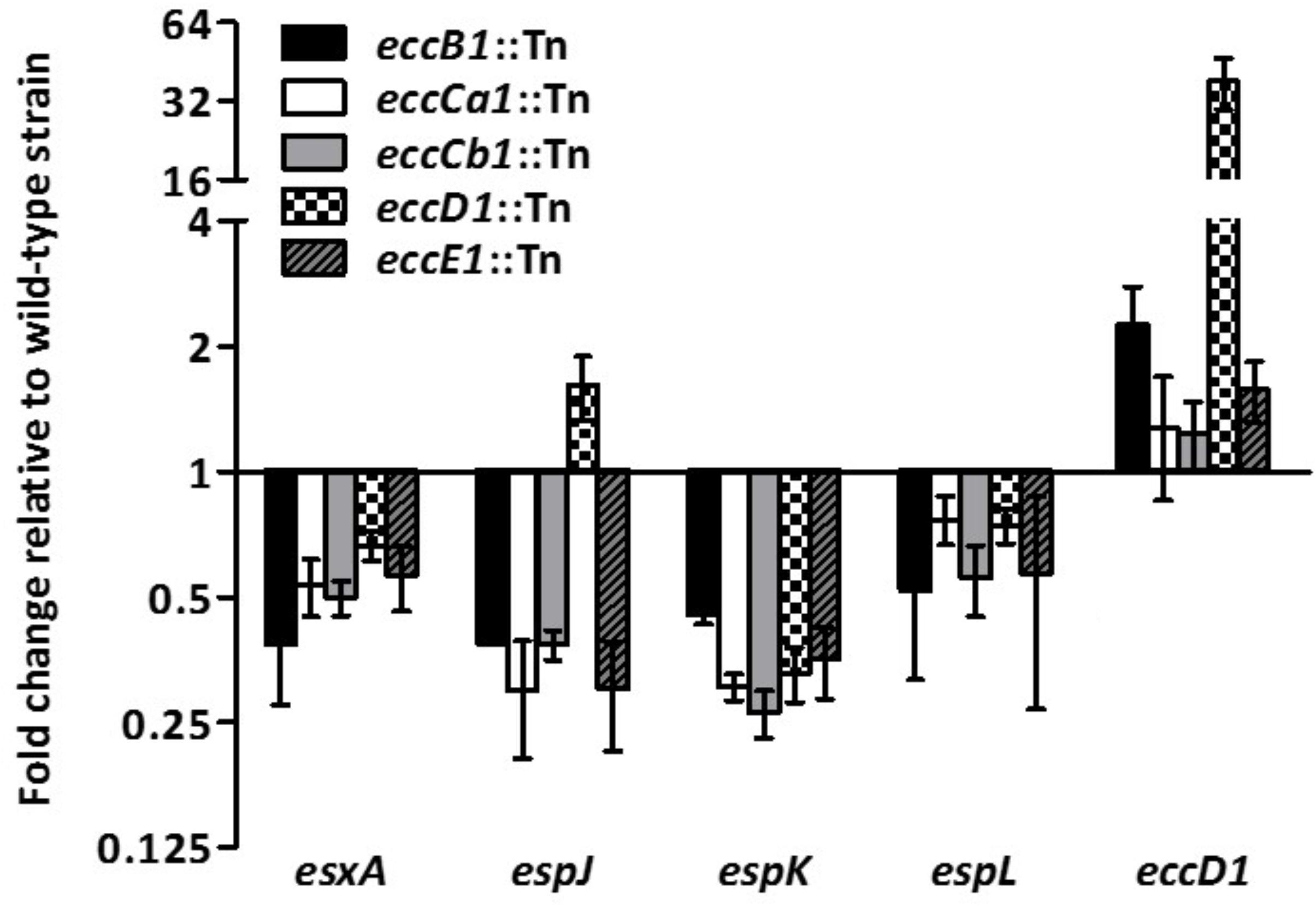
*Esx-1* Transposon Mutants have Similar Gene Transcription Profiles. Gene expression levels for *M. marinum eccB_1_*, *eccCa_1_*, *eccCb_1_*, *eccD_1_* and *eccE_1_* transposon mutants as measured by qRT-PCR. All strains were grown in 7H9 culture medium, and gene expression levels were compared to those of the wild-type strain E11. Values represent mean ± standard error of the mean of at least three biological replicates.

### ESX-1 substrate gene transcription is reduced by a regulatory mechanism

We next sought to determine the molecular mechanism underlying the down-regulation of specific transcripts in *esx-1* mutant strains of *M. marinum*. It is possible that the decrease in mRNA levels is due to a regulatory effect at the transcriptional level. Alternatively, mRNA derived from specific sequences may be degraded via a post-transcriptional mechanism. To investigate these possibilities, we expressed an extra copy of the *espL* gene under the control of a constitutively active promoter in the wild-type and *eccCb_1_* mutant strains of *M. marinum* and determined the *espL* gene transcript levels. We found a similar increase in *espL* transcripts in both the wild-type and *eccCb_1_* mutant strains, indicating that degradation of specific mRNA is probably not the cause of the decreased mRNA levels in the mutant strain (Figure 5A). Expression levels of the downstream gene *espK* were not affected by the introduction of *espL*. These results indicate that there is a regulatory mechanism that prevents the transcription of genes encoding ESX-1 substrates and associated proteins in the absence of a functionally active ESX-1.

**Fig 5.**
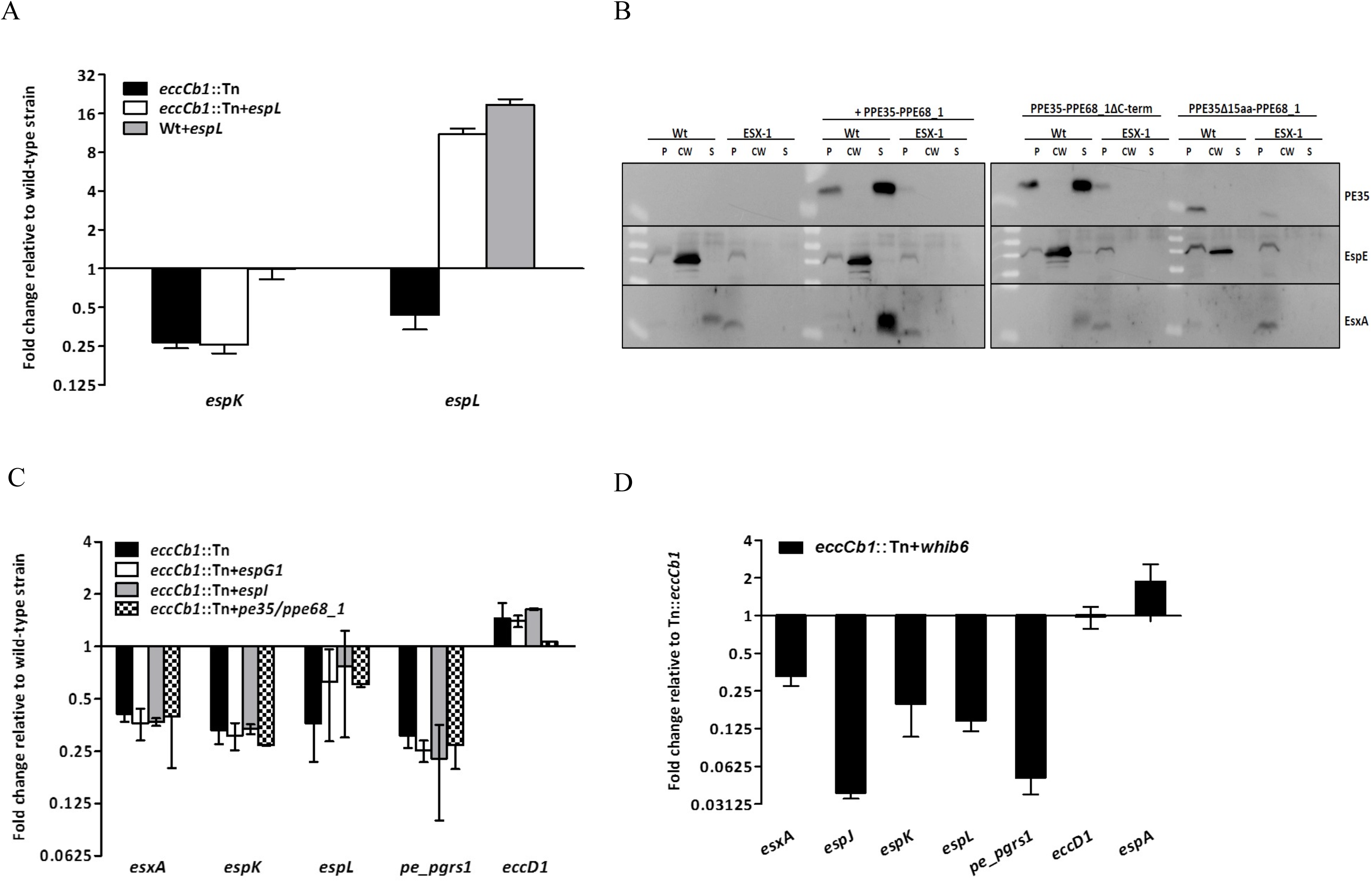
Regulation of the ESX-1 Secretion System. (A) Down-regulation of *espL* is the result of a regulatory process. A functional copy of *espL* was introduced into wild-type and *eccCb1* mutant strains of *M. marinum*, and the espK and espL gene expression levels were measured by qRT-PCR. Gene expression levels were compared to those of the wild-type strain E11. Values represent mean ± standard error of the mean of two biological replicates. (B) Introduction of PE35/PPE68_1 results in increased EsxA secretion but not in gene regulation. Pellet (p), cell wall extract (cw), and supernatant (s) fractions of the wild-type and *eccCb1* mutant strains of *M. marinum* expressing PE35/PPE68_1; PE35/PPE68, containing a C-terminal deletion of PPE68_1; or PE35/PPE68_1, containing a 15-amino-acid C-terminal deletion of PE35, were analysed for the presence of EspE, EsxA and the introduced PE35 by Western blotting. Fractions represent 0.5, 1 or 2 OD units of original culture. In all cases, PE35 contained a C-terminal HA tag. (C) EspG1, EspI and PE35/PPE68_1 do not regulate the transcription of selected esx-1-associated genes. EspG1, EspI or PE35/PPE68_1 were overexpressed in the *M. marinum* eccCb1 mutant strain, and the expression levels of *espK, espL, esxA, pe_pgrs1* and *eccD1* were measured by qRT-PCR. Gene expression levels were compared to those of the wild-type strain E11. Values represent mean ± standard error of the mean of at least two biological replicates. (D) WhiB6 is involved in transcriptional regulation of ESX-1 substrates and associated genes. The *whib6* gene was overexpressed in the *M. marinum eccCb1* mutant strain, and transcript levels of *espK*, *espL*, *esxA*, *pe_pgrs1* and *eccD1* were measured by qRT-PCR. Gene expression levels were compared to those of the *eccCb1* mutant strain. Values represent mean ± standard error of the mean of two biological replicates.

### PE35 and PPE68 play an important role in ESX-1 secretion but not in gene regulation

Previously, PE35, which is located within the *esx-1* gene cluster, has been implicated in the regulation of *esxA/esxB* gene expression in *M. tuberculosis* [34]. In contrast to this proposed function, the PE35/PPE68_1 protein pair in *M. marinum* is secreted via ESX-1 [35,36]. To determine whether PE35 plays a role in the regulation of ESX-1 substrates, we overexpressed the *pe35/ppe68_1* operon in *M. marinum*. Interestingly, although there was no effect on gene transcription (Figure 5C), we noticed a substantial increase in EsxA secretion in the wild-type strain (Figure 5B). This increased EsxA secretion does not seem to represent a general increase in ESX-1 secretion, as protein levels of the cell-surface-localized EspE remained similar (Figure 5B). To study this effect in more detail, we introduced PE35 with a truncated version of PPE68_1 that contained only the PPE domain and was devoid of the C-terminal portion. Although the introduced PE35 protein was expressed and secreted efficiently by ESX-1 (Figure 5B), the levels of secreted EsxA were not increased, indicating that the C-terminal portion of PPE68_1 plays a role in EsxA secretion. To determine whether secretion of the PE35/PPE68_1 protein pair itself was important for this process, we also determined the effect of removal of the last 15 amino acids of the PE protein, which contained the general secretion signal. This small deletion not only abolished the secretion of the introduced PE35 protein but also abolished EsxA secretion completely, despite the presence of an intact chromosomal copy of the *pe35/ppe68_1* operon (Figure 5B). This result suggests that the truncated form of PE35 somehow interferes with EsxA secretion. Together these data show that, although PE35 and PPE68_1 do not seem to regulate the transcription of genes encoding ESX-1 substrates, these proteins have a strong effect on EsxA, as previously observed [34].

### Increasing EspI and EspG_1_ levels does not lead to altered esx-1 gene expression

A second candidate protein that might regulate gene expression levels of ESX-1 substrates is EspI. The gene encoding this esx-1-secretion-associated protein of unknown function is located within the *esx-1* region and is down-regulated in *esx-1* mutants of both *M. marinum* and *M. tuberculosis* (Figure 2). In contrast to the other Esp proteins, EspI contains a putative nucleotide-binding domain. However, when we overexpressed this protein, we did not observe a change in the down-regulation of esx-1-associated gene transcription in the *M. marinum eccCb1* transposon mutant, suggesting that EspI does not regulate this process (Figure 5C). We next focused on EspG1 as a candidate *esx-1* gene regulator. EspG1, which is a cytosolic protein that is not part of the membrane-bound secretion machinery, has recently been shown to interact specifically with PE35/PPE68_1 in *M. marinum* [35]. It is conceivable that EspG_1_ might function as a sensor that measures protein levels of intracellular ESX-1 substrates. When substrate levels are low, unbound EspG1 may act as a signal for the induction of gene expression. In the absence of a functional ESX-1 system, accumulated PE35/PPE68_1 or other substrates may bind to EspG_1_, leading to reduced transcription of esx-1-associated genes. To investigate the effect of EspG_1_ on esx-1-associated gene expression and protein levels, we increased EspG1 levels by overexpressing the *espG1* gene in wild-type and ESX-1-deficient *M. marinum*. This overexpression did not result in altered gene transcription (Figure 5C) or ESX-1 protein secretion (data not shown). Together, our data show that EspI and EspG_1_ do not appear to play key roles in esx-1-associated gene regulation.

### WhiB6 plays a role in the transcription of ESX-1 substrates

In addition to *espI*, another gene encoding a putative regulatory protein was down-regulated in *esx-1* mutant strains of both *M. marinum* and *M. tuberculosis*, namely, *whiB6* (Figure 2). WhiB proteins are actinobacteria-specific regulators that contain iron-sulfur clusters and are thought to act as redox-sensing transcription factors that can cause both gene activation and repression [37]. WhiB6 was suggested to be involved in the regulation of EsxA secretion [38], and subsequent studies have confirmed this suggestion [39–41]. To determine whether Whib6 had an effect on the expression levels of *esx-1* associated genes, we overexpressed this protein in the ESX-1-deficient *M. marinum eccCb_1_* transposon mutant strain. We found that particularly those genes that were already down-regulated in the mutant strain, such as *esxA* and *espK*, showed even further transcriptional inhibition when *whib6* levels were increased (Figure 5D). Furthermore, expression of *eccD1* was unaltered by *whib6* overexpression, indicating that *whib6* is involved in the transcription of ESX-1 substrates and associated genes but not of the components of the ESX-1 system. Surprisingly, *whiB6* is one of the genes that is down-regulated upon abrogation of ESX-1-mediated protein secretion. WhiB6 may be activated when the ESX-1 machinery is blocked and represses genes encoding ESX-1 substrates as well as its own gene. Together, our data suggests that ESX-1 regulation is even more complex than previously thought.

### WhiB6 is required for the regulation of the ESX-1 system

To determine whether WhiB6 is required for ESX-1 regulation, we constructed a deletion mutant of *whiB6* both in the *M. marinum* WT strain and in the *esx-1* mutant background (*M. marinum M^USA^ -ΔwhiB6*–and *M. marinum M^VU^ -ΔwhiB6*). First, we analyzed the effect of this mutation on all genes, except for the genes of the *esx-1* locus. This analysis identified 34 genes (p<0.05) that were significantly downregulated in the *esx-1* mutant strains (Figure 6A). Complementation of both mutants with the *whiB6* gene on a mycobacterial shuttle plasmid reversed the upregulation of these genes and generally resulted in decreased expression levels (Figure 6A, B). As expected, several genes that are associated with oxidative stress (*ahpC*, *ahpD*, *rebU*) were found in the differently expressed gene pool. Also, the enrichment analysis of the associated Gene Ontology terms for the differently expressed genes (*dnaB*, *dinP*) reveal that WhiB6 may also regulate DNA replication or repair through regulating DNA-directed DNA polymerase and DNA helicase (Figure S7). Another noteworthy gene affected by *whiB6* deletion is *iniA*, which is associated with cell wall stress induced by specific antibiotics. Interestingly, *whiB7* is within the *whiB6-active* gene set, which implies that WhiB7 is active by, or works with WhiB6. Other than the *whiB6*-active gene set, 13 genes, which are involved in iron-sulfur cluster binding, cellular lipid metabolic processes are downregulated.

**Fig 6.**
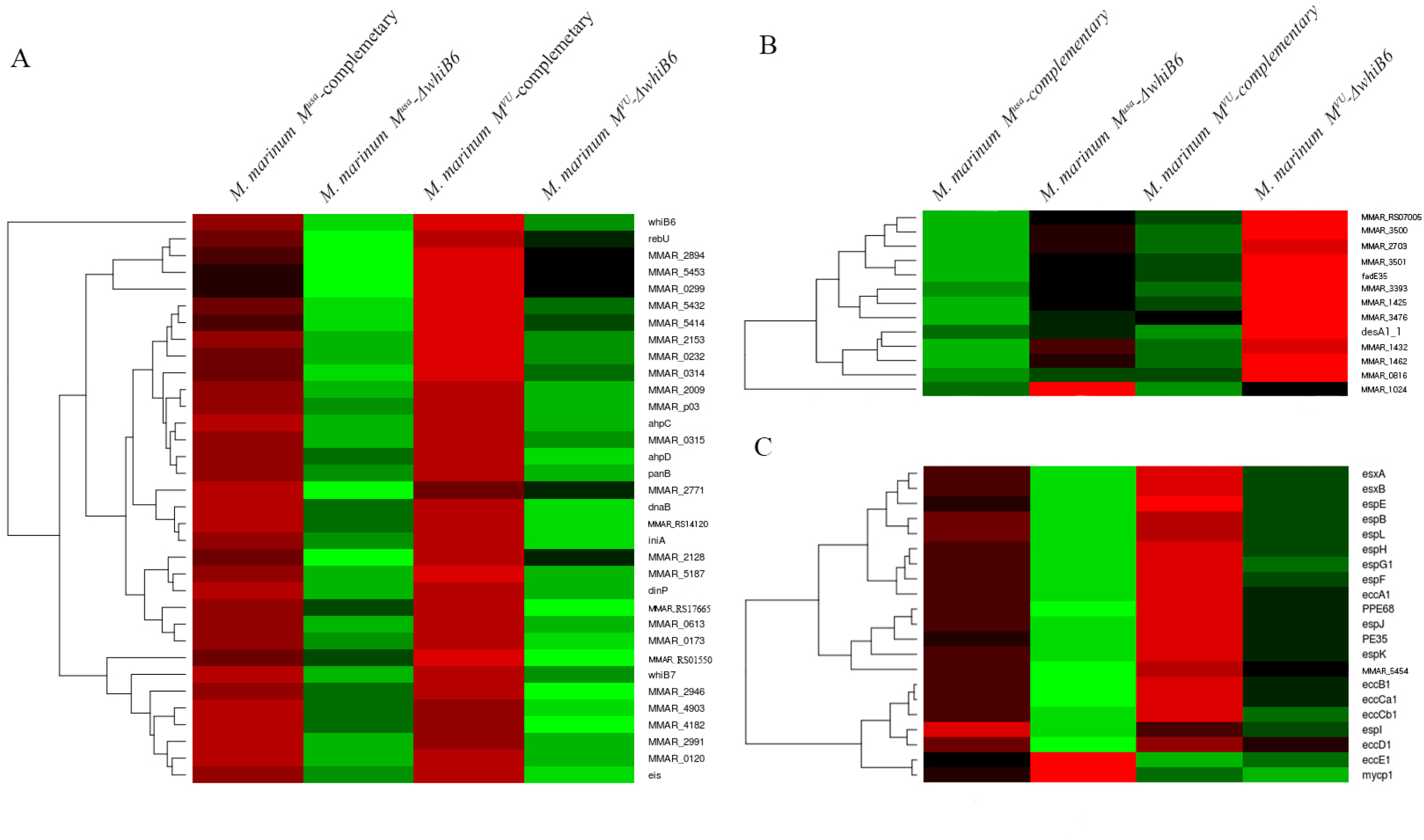
Gene Expression Profiles (Log2 Fold Change) of the Complementary Strains (*M. marinum M^USA^*-Complementary and *M. marinum M^VU^*-complementary) and Knock-Out Strains (*M. marinum M^USA^ -ΔwhiB6* and *M. marinum M^VU^ -ΔwhiB6*) Compared with those of the Corresponding Control Strains (*M. marinum M^USA^*-Empty Vector Strain and *M. marinum M^VU^-empty* vector). The green colour represents up-regulate genes and red colour represents down regulate genes compared with the control strains. The heat map of expression of the whiB6-activated gene set is shown in (A); Expression of the *esx-1* locus is shown in (B); and (C) shows the WhiB6-repressed gene set.

Remarkably, these genes are almost exclusively downregulated in the *whiB6 esx-1* double mutant, reinforcing a functional link between WhiB6 and the ESX-1 system. However, many of the downregulated genes are encoding hypothetical proteins and hence needed to be further characterized.

Separately, we analyzed the effect of the *whiB6* deletion on all *esx-1* genes. In line with our previous results, overexpression of *whiB6* in the *esx-1* mutant resulted in downregulation of many *esx-1* genes (Figure 6C), whereas deletion of *whiB6* did not have a strong affect in the *esx-1* mutant background. The effect was the opposite for the mutant with a functional ESX-1 system, there the whiB6 deletion had a strong positive effect on transcription of *esx-1* genes. (Figure 6C) and also this effect could be complemented. Only the structural *eccE1* and *mycP1* genes behaved differently. There are six genes that show the same pattern of up- or down-regulation as most of the *esx-1* genes in the two different *whiB6* mutants and the complemented strains. Of these six genes, 4 encode putative ESX-1 substrates or ESX-1 chaperones, *i.e*. MMAR_2894 (PE34-like protein), MMAR0299 (PE_PGRS1), MMAR5414 ((EspA-like) and MMAR5432 (EspD-like). Together, these experiments show a strong linkage between WhiB6 and the regulation of different *esx-1* genes in response to secretion activity.

## Discussion

In this study, we determined the transcriptomes of the *M. marinum* E11 wild-type and the double-auxotrophic *M. tuberculosis* mc^2^6020 mutant strains and compared these transcriptomes with those of the corresponding isogenic *esx-1* mutants. We found that during growth in 7H9 culture medium, genes encoding ESX-1 substrates, such as EsxA and other ESX-1-associated proteins, were down-regulated in the mutant strains, whereas the transcription of genes encoding several structural components of the ESX-1 system remained unaffected. This specific decrease in transcription might function as a mechanism to avoid toxic accumulation of ESX-1 substrates. Interestingly, a similar decrease in substrate production has been shown for the ESX-5 secretion system, where the PE_PGRS substrates do not accumulate intracellularly when secretion is blocked [42,43]. However, for these PE_PGRS substrates, regulation was shown to occur post-transcriptionally [43], implying that a different mechanism is involved.

The most prominent change in gene expression that was observed upon host cell infection by the *M. marinum esx-1* mutant strain was the increase in transcription of the *espA* operon. The specific and pronounced transcriptional increase in the expression of this operon, and not of any other *esx-1* associated gene, indicates that transcription of the *espA* operon is regulated independently of the other substrates during infection. Previously, it has been shown that the *espA* operon is regulated by different transcription and regulation factors, including EspR, MprAB and PhoPR [20,44,45]. Our new finding also suggests that EspA, EspC and EspD are vital for the bacteria during the early phase of infection. Since ESX-1 has been shown to be responsible for mycobacterial escape from the phagosome, which occurs within the first few hours of infection with *M. marinum* [6], the proteins produced by the *espA* operon may play an important role in this process. Consequently, the avirulent phenotype of ESX-1-deficient mycobacteria might be partly attributable to the inability to secrete EspA and/or EspC early in infection.

To determine the mechanism via which ESX-1 substrate regulation is mediated, we overexpressed proteins that may have a regulatory function. Overexpression of the *esx-1-*encoded EspI and EspG_1_ proteins did not have an effect on the reduced transcription of ESX-1 substrates in ESX-1-deficient *M. marinum*. The putative regulatory protein WhiB6, however, did affect the transcription of these genes. While the transcript levels of *whib6* itself were decreased in *esx-1* mutants of *M. marinum* and *M. tuberculosis*, increasing WhiB6 levels by overexpression resulted in a further decrease in transcription of the ESX-1 substrate in ESX-1-deficient *M. marinum*. This result clearly indicates that WhiB6 is involved in ESX-1-associated gene regulation, as previously suggested [41]. Indeed, there is accumulating evidence that WhiB proteins function as transcription factors that may play a role in survival within the host (reviewed in [46]). Recently, other groups have also presented evidence supporting a role for WhiB6 in the regulation of the transcription of *esx-1* genes [39–41].

A remarkable finding in this study was that overproduction of PE35/PPE68_1 resulted in a large increase in EsxA secretion. Previously, deletion of *M. tuberculosis* PE35 had already been shown to abolish *esxA* transcription and secretion of the corresponding gene product [34]. Here, we found that EsxA and PE35 secretion are linked, as an increase in PE35 secretion resulted in a concomitant increase in EsxA secretion. The C terminus of PPE68_1 is required for this effect, indicating that this is a specific process, which is consistent with the fact that cell-surface localization of another ESX-1 substrate, namely, EspE, is unaffected by overproduction of PE35/PPE68_1. It is possible that the PPE68 proteins serve as chaperones to escort EsxA outside the bacterium, or these proteins, may be part of the secretion apparatus, making the secretion of specific substrates highly efficient.

During *M. marinum* infection of human macrophages, we found that transcription of many *pe_pgrs* and *ppe* family genes was strongly down-regulated when ESX-1 function was abrogated. As many as 50% of all genes with decreased transcript levels in the *esx-1* mutant strain belongs to one of these gene families (Table S4). Notably, in the wild-type strain, transcription of the *pe_pgrs* and *ppe* genes was decreased during infection in comparison to the levels observed during growth in 7H9 medium (Table S5). As part of an adaptive response to the macrophage environment, expression of these cell-wall-localized proteins may be down-regulated in order to evade immune recognition or to reduce cell permeability [47]. The fact that in the absence of a functional ESX-1 secretion system these genes are even further down-regulated suggests that there are functional links or shared transcriptional pathways between ESX-1 and (some of the) PE_PGRS and PPE proteins, which are generally ESX-5 substrates [48].

Taken together, our results show that transcription of the *espA* locus plays an important role in ESX-1 mediated processes during the first hours of infection. Furthermore, we established a functional link between PE35 and EsxA secretion and provided evidence of a regulatory role of WhiB6 in the transcription of ESX-1 substrates and associated genes.

## Materials and Methods

### Bacterial strains and growth conditions

The *esx-1* mutants of the *M. marinum* E11 wild-type strain used in this study contain transposon insertions in *eccB_1_, eccCa_1_, eccCb_1_, eccD_1_* and *eccE_1_* [49]. For *M. tuberculosis*, the attenuated double-deletion strains mc^2^6020 and mc^2^6030 of H37Rv were used, with deletions of *lysA* and *panCD* and of *RD1* and *panCD*, respectively [24,50]. Bacterial strains were grown with shaking at 30°C (*M. marinum*) or 37°C (*M. tuberculosis*) in Middlebrook 7H9 culture medium supplemented with 10% ADC (albumin-dextrose-catalase, BD Biosciences) and 0.05% Tween-80. Culture medium containing the auxotrophic *M. tuberculosis* deletion strains was supplemented with 50 μg/ml pantothenic acid and, for mc^2^6020, 100 μg/ml L-lysine.

### Infection of human macrophages

THP-1 monocytes were cultured at 37°C in 5% CO2 in RPMI-1640 with GlutaMAX-1 (Gibco) supplemented with 10% FBS, 100 μg/ml streptomycin and 100 U/ml penicillin. Cells were seeded at a density of 3 × 10^7^ cells per T175 flask and differentiated into macrophages by 48 hours of incubation with 25 ng/ml PMA (Sigma-Aldrich). Then, 1.8 × 10^8^ THP-1 cells were infected with *M. marinum* at a multiplicity of infection (MOI) of 20 for 2 hours, after which the cells were washed with PBS to remove extracellular bacteria. After 4 additional hours of infection at 33°C, the THP-1 cells were lysed with 1% Triton X-100. After a low-speed centrifugation step to remove cellular debris, mycobacteria were pelleted, after which RNA was extracted as described in the following section.

### Genome sequence

We sequenced the *M. marinum* E11 strain with PacBio RSII single-molecule real-time (SMRT) sequencing technology [51]. The raw reads were assembled into two pieces (the core and the plasmid) with HGAP assembler [52] using the default parameters. The sequence was improved with iCORN2 [53] with three iterations, correcting 20 single base pair errors and 61 insertions and deletions. To transfer the annotation from the current reference, we used RATT [54] with the PacBio parameter. Gene models around gaps were manually improved on the new sequence. The updated genome annotation was resubmitted under the same accession numbers (HG917972 for the *M. marinum E11* main chromosome genome and HG917973 for the *M. marinum E11* pRAW plasmid; complete sequences).

### RNA extraction and qRT-PCR

*M. marinum* and *M. tuberculosis* cultures were pelleted and bead beated in 1 ml of TRIzol (Invitrogen) with 0.1-mm zirconia/silica beads (BioSpec Products). After centrifugation, supernatants were extracted with chloroform, and RNA was precipitated with isopropanol. RNA pellets were washed with 80% ethanol and dissolved in RNAse-free water. Contaminant DNA was removed by incubation with DNAse I (Fermentas). For RT-PCR, cDNA was generated using a SuperScript VILO cDNA Synthesis Kit (Invitrogen). An equivalent of 5 ng of RNA was used in the quantitative PCRs. qRT-PCR was performed using SYBR GreenER (Invitrogen) and a LightCycler 480 (Roche) instrument. Transcript levels were normalized to the levels of the housekeeping gene *sigA* [55] using ΔΔCt analysis. All primer sequences used for qRT-PCR are listed in Table S8.

### RNA preparation for Illumina sequencing

Total RNA was extracted with TRIzol (Invitrogen) and then purified on RNeasy spin columns (Qiagen) according to the manufacturer’s instructions. RNA integrity (RNA integrity score ≥ 6.8) and quantity were determined on an Agilent 2100 Bioanalyzer (Agilent; Palo Alto, CA, USA). As ribosomal RNA constitutes a vast majority of the extracted RNA population, depletion of these molecules via RiboMinus-based rRNA depletion was conducted. For mRNA enrichment, Invitrogen’s RiboMinus Transcriptome Isolation Kit, bacteria was used according to manufacturer’s instructions. Briefly, 2 μg of total RNA samples was hybridized with prokaryotic rRNA-sequence-specific 5’-biotin-labelled oligonucleotide probes to selectively deplete large rRNA molecules from total RNA. Then, these rRNA-hybridized, biotinylated probes were removed from the sample with streptavidin-coated magnetic beads. The resulting RNA sample was concentrated using the RiboMinus concentrate module according to the manufacturer’s protocol. The final RiboMinus RNA sample was subjected to thermal mRNA fragmentation using the Elute, Prime, Fragment Mix from the Illumina TruSeq RNA Sample Preparation Kit v2 (Low-Throughput protocol). The fragmented mRNA samples were subjected to cDNA synthesis using the Illumina TruSeq RNA Sample Preparation Kit (low-throughput protocol) according to the manufacturer’s protocol. Briefly, cDNA was synthesized from enriched and fragmented RNA using SuperScript III reverse transcriptase (Invitrogen) and the SRA RT primer (Illumina). The cDNA was further converted into double-stranded DNA using the reagents supplied in the kit, and the resulting dsDNA was used for library preparation. To this end, cDNA fragments were end-repaired and phosphorylated, followed by adenylation of 3 ends and adapter ligation. Twelve cycles of PCR amplification were then performed, and the library was finally purified with AMPure beads (Beckman Coulter) as per the manufacturer’s instructions. A small aliquot (1 μl) was analysed on an Invitrogen Qubit and an Agilent Bioanalyzer. The bar-coded cDNA libraries were pooled at equal concentrations before sequencing on an Illumina HiSeq2000 using the TruSeq SR Cluster Generation Kit v3 and TruSeq SBS Kit v3. Data were processed with Illumina Pipeline software v1.82.

### RNA-seq analysis

The Illumina reads were mapped with SMALT (http://www.sanger.ac.uk/science/tools/smalt-0) (default parameters) against the new PacBio reference. From the read count, which was obtained with bedtools ([56], parameter multicov, with -D to include duplicates and -q 5 to exclude repetitive mapping reads), we performed a differential expression analysis with DESeq [57] using default parameters.

### Plasmid construction

The *E*. coli-mycobacterial shuttle vector pSMT3 was used for the construction of all plasmids. To overexpress PE35-PPE68_1 (MMARE11_01740-MMARE11_01750), we used a previously described plasmid [13]. For construction of the plasmid containing *espG_1_*, this gene was amplified from the *M. marinum* E11 genome by PCR using primers containing NheI and EcoRV restriction sites and a 3’ HA epitope. The resulting PCR product and empty pSMT3 were digested with NheI and EcoRV followed by ligation of *espG_1_* into the vector by T4 ligase (Fermentas). For construction of the plasmid containing *whib6*, this gene was amplified from the *M. marinum* E11 genome by PCR using primers containing NheI and BamHI restriction sites. For the other construct, *espI* was amplified from the *M. marinum* E11 genome by PCR using primers containing NheI and BglII restriction sites. The PCR product was digested with NheI and BamHI. Empty pSMT3 was digested with NheI and BamHI, after which the PCR product was ligated into the vector. All plasmids were introduced into the *M. marinum* wild-type E11 and isogenic *eccCb_1_* mutant strains by electroporation. All primer sequences are listed in Table S8.

### Analysis of protein expression and secretion

*M. marinum* cultures were grown to mid-logarithmic phase in 7H9 culture medium supplemented with 0.2% glycerol and 0.2% dextrose. Bacteria were pelleted, washed in PBS and incubated in 0.5% Genapol X-080 (Sigma-Aldrich) for 30 minutes to extract cell wall proteins. Genapol X-080-treated *M. marinum* cells were disrupted by sonication. Secreted proteins were precipitated from the culture supernatant by 10% trichloroacetic acid (TCA, Sigma-Aldrich). Proteins were separated according to molecular weight on 15% SDS-PAGE gels and subsequently transferred to nitrocellulose membranes (Amersham Hybond ECL, GE Healthcare Life Sciences). Immunostaining was performed with mouse monoclonal antibodies directed against the HA epitope (HA. 11, Covance), EsxA (Hyb76-8), or rabbit polyclonal sera recognizing EspE [58].

### LC-MS analysis

Peptide preparation from the *M. marinum* E11 and isogenic *esx-1* mutant strains was performed as previously described [59]. Approximately 100-μg protein digests of each sample were labelled with 4plex iTRAQ reagents (Applied Biosystems). The combined iTRAQ-labelled samples were fractionated using strong cation exchange chromatography. The eluted fractions were dried and desalted using a Sep-Pak C-18 SPE cartridge (Waters, Milford, MA, USA). LC-MS analysis as well as MS data processing was carried out following our published procedure [60]. Briefly, each fraction was analysed three times using an LTQ-Orbitrap Velos (Thermo Scientific). The MS spectra were recorded in the Orbitrap, whereas the MS2 spectra were recorded in the c-TRAP for HCD fragmentation and in the LTQ for the CID fragmentation. Both HCD and CID spectra were extracted separately using Proteome Discoverer software and processed by an in-house script before a Mascot search against the *M. marinum* E11 proteome. The Mascot results (.dat file) were processed by Scaffold software for validation of protein identification and quantitative assessment. For protein identification, local false positive rates (FDR) were maintained below 1% for both protein and peptide identification (0.91% and 0.9% for peptides and proteins, respectively, for this dataset). Protein quantitation was processed using Scaffold Q+, which is based on the i-Tracker algorithm [61]. The iTRAQ quantitation using HCD is highly accurate, and a change of more than 2-fold was considered significant differential expression in this study.

## Accession codes

Sequencing reads have been submitted to the EMBL-EBI European Nucleotide Archive (ENA) Sequence Read Archive (SRA) under the study accession no. PRJEB8560.

## Acknowledgements

We thank Astrid van der Sar and Esther Stoop for providing the *M. marinum E11* ESX-1 mutants. Work in AP’s laboratory is supported by the KAUST faculty baseline fund (BAS/1/1020-01-01). The authors thank members of the Bioscience Core Lab (BCL) at KAUST for sequencing the RNA-seq libraries on the Illumina Hiseq platform and for running protein samples through the quantitative proteomics workflow with the LTQ-Orbitrap Velos instrument (Thermo Scientific).

## Supplementary table and figure legends

**Table S1.**

Complete list of genes for which the expression levels changed significantly in the *M. marinum eccCb_1_* transposon mutant compared to the levels in the isogenic wild-type strain E11 during growth in 7H9 culture medium. P<0.05.

**Table S2.**

Complete list of proteins for which the expression levels changed in *M. marinum eccCb_1_* transposon mutant compared to the levels in the isogenic wild-type strain E11 during growth in 7H9 culture medium. Proteins with greater than 2-fold change were considered significantly differentially expressed.

**Table S3.**

Complete list of genes for which the expression levels changed significantly (p<0.05) in the auxotrophic *M. tuberculosis* RD1 deletion mutant strain mc^2^6030 compared to the levels in the isogenic control strain mc^2^6020 during growth in 7H9 culture medium.

**Table S4.**

Complete list of genes for which the expression levels changed significantly (p<0.05) in the *M. marinum eccCb_1_* transposon mutant strain compared to the levels in the wild-type strain E11 during infection of human THP-1 macrophages.

**Table S5.**

Complete list of genes for which the expression levels changed significantly (p<0.05) in the *M. marinum* wild-type strain during infection of macrophages compared to the levels during growth in 7H9 culture medium.

**Table S6.**

Complete list of genes for which expression levels changed significantly (p<0.05) in the *M. marinum eccCb_1_* transposon mutant strain during infection of macrophages compared to the levels in the wild-type strain E11 during growth in 7H9 culture medium.

**Table S7.**

Complete list of genes for which the expression levels changed significantly (p<0.05) in the *M. marinum eccCb_1_* transposon mutant strain during infection of macrophages compared to the levels during growth in 7H9 culture medium.

**Table S8.**

Primers used in this study. Restriction sites are shown in bold.

**Figure S1**

Euclidean distance matrices of RNA-seq transcriptome data showing clustering of *M. marinum* wild-type (E11) and *eccCb_1_* transposon mutant (ESX-1) strains grown in culture medium (three biological replicates) or during infection of THP-1 cells (indicated as ‘int’).

**Figure S2**

Principal component analysis (PCA) of biological replicates of proteome data showing clustering of *M. marinum* wild-type (E11) and *eccCb_1_* transposon mutant (ESX-1) strains. PCA mapping showed clustering of biological replicates of the E11 wild-type and *esx-1* mutant strains.

**Figure S3**

Correlation between protein and mRNA expression of the *M. marinum eccCb_1_* transposon mutant and the isogenic wild-type strain E11 during growth in 7H9 culture medium. (A) Scatterplot of the relationship between differentially expressed genes quantified in both data sets. (B-F) Scatterplots for protein and gene transcript expression classified by functional categories. Scatterplots display the rectilinear equation and the Pearson correlation coefficient (*R^2^*).

**Figure S4**

Functional categories of genes that are significantly changed in the transcriptome and proteome of the *M. marinum eccCb_1_* transposon mutant compared to the isogenic wild-type strain E11 during growth in 7H9 culture medium. Genes exhibiting differential expression at the RNA and protein levels were grouped according to the MarinoList classification (http://mycobrowser.epfl.ch/marinolist.html).

**Figure S5**

Most differentially expressed genes of the *M. marinum eccCb_1_* transposon mutant compared to the isogenic wild-type strain E11 during infection of primary macrophages, grouped into broad functional categories. Within each group, genes are ranked in ascending order by *P*-value. (A). Top 100 annotated genes from the *M. marinum E11* strain that were the most differentially expressed in the *M. marinum* wild-type strain E11 during infection of primary macrophages. Bar chart of log2-fold changes for individual genes (tags, left). (B). Top 100 annotated genes from the *M. marinum E11* strain that were the most differentially expressed in the *M. marinum eccCb_1_* transposon mutant compared to the isogenic wild-type strain E11 during infection of primary macrophages (tags, left). Bar chart of log2-fold changes for individual genes.

**Figure S6:**

Regulation of genes encoding proteins predicted to be involved in metabolic adaptation, energy metabolism and transcriptional regulatory processes in the *M. marinum eccCb_1_* transposon mutant grown in 7H9 culture medium as well as in the wild-type and *eccCb_1_* transposon mutant strains during infection in human THP-1 macrophages (indicated as ‘int’) compared to the that in the wild-type strain E11 during growth in 7H9 culture medium. (A) *Catabolism of fatty acids*. Genes were selected based on their annotation and ordered based on expression. (B) Energy generation and NAD+ regeneration. Genes were selected based on their annotation and ordered based on expression. (C) Genes of the *kstR* regulon, which are required for uptake and metabolism of cholesterol (1, 2). (D) Transcriptional regulation. Genes were selected based on their annotation and ordered based on expression.

**Figure S7**

The enriched Gene Ontology (GO) terms of the gene set activated (excluding the *esx-1* locus genes) or repressed by WhiB6. The molecular function GO terms are in red, while the biological process terms are in blue.

